# Fundamental Law of Memory Recall

**DOI:** 10.1101/510750

**Authors:** Michelangelo Naim, Mikhail Katkov, Sandro Romani, Misha Tsodyks

## Abstract

Free recall of random lists of words is a standard paradigm used to probe human memory. We proposed an associative search process that can be reduced to a deterministic walk on random graphs defined by the structure of memory representations. The corresponding graph model is different from the ones considered in the past but still can be solved analytically, resulting in a novel parameter-free prediction for the average number of memory items recalled (*RC*) out of *M* items in memory: 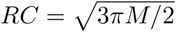. This prediction was verified with a specially designed experimental protocol combining large-scale crowd-sourced free recall and recognition experiments with randomly assembled lists of words or common facts. Our theoretical and experimental results indicate that memory recall operates according to a stereotyped search process common to all people.

---

Human memory capacity for storing information is remarkable, but recalling a collection of unrelated events is challenging. To understand human memory one needs to understand both the ability to acquire vast amounts of information and at the same time the limited ability to recall random material. The standard experimental paradigm to address the later question is free recall (e.g. see Kahana 2012). Typical experiments involve recalling randomly assembled lists of words in an arbitrary order after a brief exposure. In these experiments, when the presented list becomes longer, the average number of recalled words grows but in a sublinear way, such that the fraction of words recalled steadily decreases (Binet and Henri 1894, Standing 1973, Murray et al. 1976). The exact mathematical form of this relation is controversial and was found to depend on the details of experimental procedures, such as presentation rate (Waugh 1967). In some studies, recall performance was argued to exhibit a power-law relation to the number of presented words (Murray et al. 1976), but parameters of this relation were extremely variable across different experimental conditions.

In our recent publications (Romani et al. 2013, Katkov et al. 2017) we proposed to treat information recall from memory as a deterministic step-by-step associative recall process. We showed with simulations (Katkov et al. 2017) that the model qualitatively accounts for a power-law behavior of recall and provides a good quantitative fit to published experimental results for a particular number of presented words (16) (Healey et al. 2014). Here we show that this model can be solved analytically and confirmed with a specially designed experimental protocol.

The recall process proposed in (Romani et al. 2013, Katkov et al. 2017) is based on two principles:

- Memory items are represented in the brain by overlapping random sparse neuronal ensembles in dedicated memory networks;
- The next item to be recalled is the one with a largest overlap to the current one, excluding the item that was recalled on the previous step.

The first principle is a common element of most neural network models of memory (see e.g. Hopfield [1982], Tsodyks and Feigel’man [1988]), while the second one is inspired by “Search of Associative Memory” (SAM, elaborated later). More specifically, item representations are chosen as random binary {0,1} vectors where each element of the vector chosen to be 1 with small probability *f* ≪ 1 independently of other elements. Overlaps are defined as scalar products between these representations. The model is illustrated in Fig. 1 (more details in Supplemental Material), where the matrix of overlaps (‘similarity matrix’, or SM) between 16 memory representations is shown in Fig. 1a. Fig. 1b is a graph that shows the transitions between memory items induced by the SM. When the first item is recalled (say the 1st one in the list), the corresponding row of the matrix, which includes the overlaps of this item with all the others, is searched for the maximal element (14^th^ element in this case), and hence the 14^th^ item is recalled next. This process continues according to the above rule (black arrows), unless it points to an item that was just recalled in the previous step, in which case the next largest overlap is searched (red arrows). After a certain number of transitions, this process begins to cycle over already visited items. This happens either the first time a previously recalled item is reached again, or the process could make some number of transitions over previously recalled items (items 10, 14, 1 in Fig. 1b) to open up a new trajectory (items 13, 5, 11, 12) until finally converging to a cycle. After the cycle is reached, no new items can be recalled.

**Figure 1.**
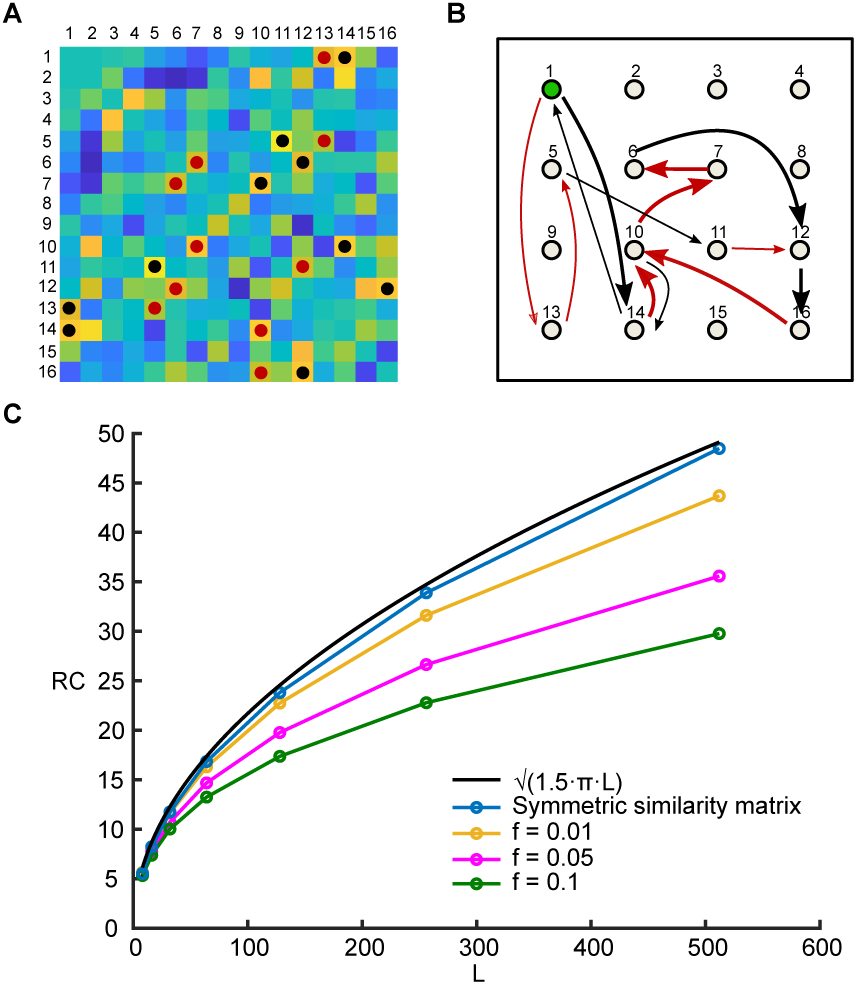
Associative search model of free recall. (**A**) SM (similarity matrix) for a list of 16 items (schematic). For each recalled item, the maximal element in the corresponding row is marked with a black spot, while the second maximal element is marked with a red spot. (**B**) A graph with 16 nodes illustrates the items in the list. Recall trajectory begins with the first node, and proceeds to an item with the largest similarity to the current one (black arrow) or the second largest one (red arrow) if the item with the largest similarity is the one recalled just before the current one. When the process returns to the 10th item, a second sub-trajectory is opened up (shown with thinner arrows) and converges to a cycle after reaching the 12^th^ node for the second time. (**C**) Comparison between simulations with random symmetric similarity matrix (blue line) and SM defined by random sparse ensembles with sparsity *f* = 0.01 (yellow line), *f* = 0.05 (magenta line), *f* = 0.1 (green line) and *N* = 100000 number of neurons. Each point is the mean of 10000 simulations. Black line corresponds to 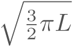.

In our previous publication (Romani et al. 2013) we showed that the average number of recalled items (recall capacity, or *RC*) scales as a power-law function of the number of items in the list, *L* with exponent that depends on sparseness parameter *f*. Here we focus on the sparse limit of this model, *f* ≪ 1, when one can neglect the correlations between different elements of the SM and replace it by a *random symmetric* matrix (see e.g. Quian Quiroga and Kreiman [2010], for biological motivation for considering a very sparse encoding). We show below that while the corresponding graph model has a history-dependent transition rule and hence is more complex than the standard family of graphs resulting from random mappings (see e.g.Harris [1960]), it can still be solved analytically in terms of the average number of items visited before converging to a cycle.

It is instructive to first consider the simpler case of a fully random asymmetric SM with independent elements. In this case, transitions between any two items are equally likely, with probability 1*/*(*L* − 1). When an item is reached for the second time the process enters into a cycle. Therefore the probability that *k* out of *L* items will be retrieved is simply

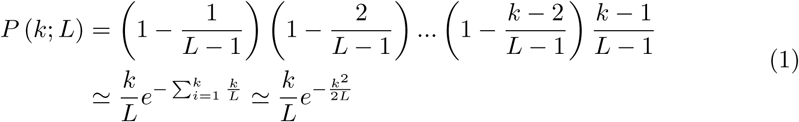

where we considered a limit of large number of items in the list (*L* ≫ 1) and assumed that *L* ≫ *k* ≫ 1, which is confirmed a posteriori below. The average number of recalled words can then be calculated as

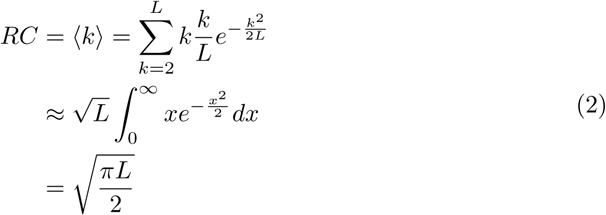

which is a well known result in random graphs literature (Harris [1960], Katz et al. [1996]).

When the SM is symmetric, as in our case, the statistics of transitions in the corresponding graph is more complicated (see Supplemental Material for more details about the derivation). In particular, the probability for a transition to one of the previously recalled items scales as 1*/*(2*L*) rather than 1*/L* as in the case of asymmetric SMs, and hence the average length of trajectory until the first return converges to 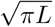. Moreover, with probability 1*/*3 the trajectory then turns towards previous items and opens up a new route until again hitting a previously recalled item, etc. Taken together, the chance that recall trajectory enters a cycle after each step asymptotically equals to 1*/*(2*L*) · 2*/*3 = 1*/*(3*L*), as opposed to 1*/L* for the fully random matrix, and hence the *RC* can be obtained by replacing *L* by 3*L* in Eq. (2):

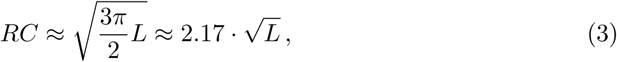

see Fig. 1c for the comparison of this analytical estimate with numerical simulations of the model. We emphasize that Eq. (3) does not have any free parameters that could be tuned to fit the experimental results. Hence, both the exponent and coefficient of this power law expression are a result of the assumed recall mechanism; in other words, this equation constitutes a true prediction regarding the asymptotic recall performance for long lists of items as opposed to earlier theoretical studies. Here we present the results of our experiments designed to test this prediction.

The universality of the above analytical expression for RC seems to contradict our everyday observations that people differ in terms of their memory effectiveness depending, e.g. on their age and experience. Moreover, it is at odds with previous experimental studies showing that performance in free recall task strongly depends on the experimental protocol, for example presentation rate during the acquisition stage (see e.g. Murdock Jr 1960, 1962, Roberts 1972, Howard and Kahana 1999, Kahana et al. 2002, Zaromb et al. 2006, Ward et al. 2010, Miller et al. 2012, Grenfell-Essam et al. 2017) and the extent of practice (Klein et al. 2005, Romani et al. 2016). Since most of the published studies only considered a limited range of list lengths, we performed free recall experiments on the Amazon Mechanical Turk^®^ platform for list lengths of 8; 16; 32; 64; 128; 256 and 512 words, and two presentation rates: 1 and 1:5 seconds per word. To avoid practice effects, each participant performed a single free recall trial with a randomly assembled list of words of a given length. The results confirm previous observations that recall performance improves as the time allotted for acquisition of each word increases, approaching the theoretical prediction of Eq. (3) from below (see Fig. 2a).

**Figure 2.**
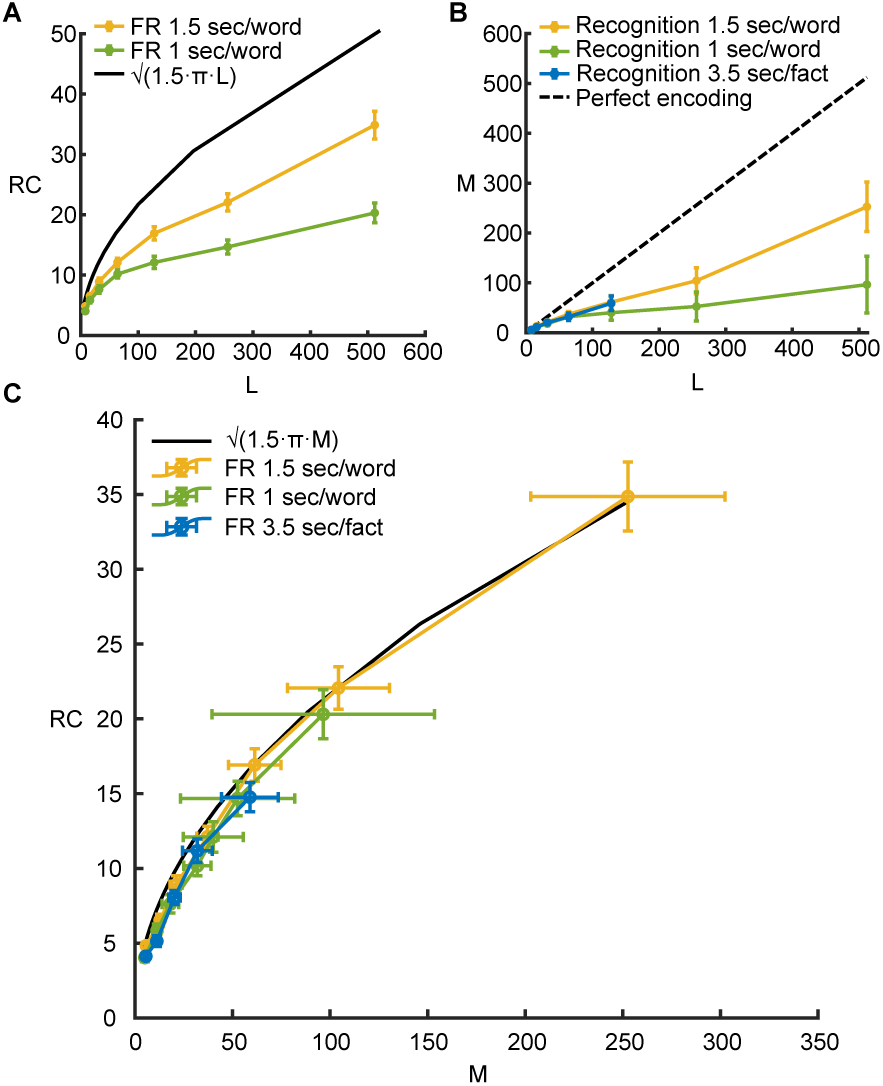
Human recall and recognition performance. (**A**) Average number of words recalled as a function of the number of words presented. Black line: Eq. (3). Yellow line: experimental results for presentation rate 1.5 sec/word. Green line: experimental results for presentation rate 1 sec/word. The error in *RC* is a standard error of the mean. (**B**) Estimated average number of acquired words/sentences for lists of different lengths. Black dashed line corresponds to perfect encoding, green line corresponds to presentation rate 1 sec/word and yellow line to presentation rate 1.5 sec/word; blue line corresponds to lists of short 8, 16, 32, 64, 128, 256 and 512 words, and two presentation rates: 1 and 1.5 seconds per word. To avoid practice effects, each participant performed a single free recall trial with a randomly assembled list of words of a given length. The results confirm previous observations that recall performance improves as the time allotted for acquisition of each word increases, approaching the theoretical prediction of Eq. (3) from below (see Fig. 2a). We reasoned that some or all of the variability in the experimentally observed RC could be due to non-perfect acquisition of words during the presentation phase of the experiment, such that some of the presented words are not encoded in memory well enough to be candidates for recall. It seems reasonable that acquisition depends on various factors, such as attention, age of participants, acquisition speed, etc. One should then correct Eq. (3) for RC, replacing the number of presented words *L* with the number of effectively acquired words *M* : sentences (see text for details). The error in *M* is computed with To test this conjecture, we designed a novel experimental protocol that bootstrap procedure (Efron and Tibshirani 1994). (**C**) Average number of words/sentences recalled as a function of the average number of acquired words. Black line: theoretical prediction, Eq. (4). Green line: experimental results for presentation rate 1 sec/word. Yellow line: experimental results for presentation rate 1.5 sec/word. Blue line: experimental results for short sentences. The error in *RC* is a standard error of the mean, while the error in *M* is computed with bootstrap procedure (see Supplemental Material for details).

We reasoned that some or all of the variability in the experimentally observed RC could be due to non-perfect acquisition of words during the presentation phase of the experiment, such that some of the presented words are not encoded in memory well enough to be candidates for recall. It seems reasonable that acquisition depends on various factors, such as attention, age of participants, acquisition speed, etc. One should then correct Eq. (3) for RC, replacing the number of presented words *L* with the number of effectively acquired words *M*:

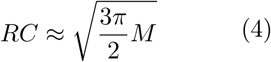

To test this conjecture, we designed a novel experimental protocol that involved performing both recall and *recognition* experiments on the same group of participants and under identical presentation conditions, in order to independently evaluate the average number of words effectively acquired and recalled upon presentation of a list of length L. Following Standing [1973], we presented participants with pairs of words, one from the list just presented (target) and one randomly chosen lure, requesting them to report which word was from the list. The average number of words acquired (*M*) was then estimated from the fraction of correctly recognized words by assuming that if a target word was effectively acquired during presentation, it will be chosen during recognition, otherwise the participant will randomly guess which of the two words is a target: *M* = *L* (2*c* − 1). Importantly, each participant performed a single recognition test, to avoid the well known effect of ‘output interference’ between subsequent recognition tests for a single list (see e.g. Criss et al. 2011).

Fig. 2b shows the estimated average number of acquired words *M* as a function of list length *L* (see Supplemental Material for details of analysis). Results confirm that acquisition improves with time allotted to presentation of each word. Standard error of the mean for the number of acquired words across participants, for each list length and each presentation speed, was estimated with a bootstrap procedure by randomly sampling a list of participants with replacement (Efron and Tibshirani 1994, see Supplemental Material).

In Fig. 2c experimentally obtained RC (yellow and green lines) is compared with the theoretical prediction of Eq. (4) (black line), where *M* is the average number of encoded words, estimated in the recognition experiment. Remarkably, agreement between the data and theoretical prediction is very good for both presentation rates, even though the number of acquired and recalled words is very different in these two conditions for each value of list length. We also performed multiple simulations of our recall algorithm (Romani et al. 2013, Katkov et al. 2017) and found that it captures the statistics of the recall performances as accessed with bootstrap analysis of the results (see Fig. S1 in Supplemental Material).

Experiments presented above, as well as the vast majority of previous recall experiments, were performed with lists of words. To test the generality of our model, we generated a set of 325 short sentences expressing common knowledge facts, such as ‘Earth is round’ or ‘Italians eat pizza’, etc. We repeated our experiments with random lists of 8, 16, 32, 64, and 128 such sentences, each presented for 3.5 seconds (see Supplementary Material for more details of the analysis). As shown in Figs. 2b and 2c, performance with lists of sentences is very close to that of words with 1.5 words per second presentation rate, albeit with some small deviations towards lower levels.

## Discussion

The results presented in this study show that the relation between the number of acquired and recalled words conforms with remarkable precision to the analytical, parameter-free expression Eq. (4), derived from a deterministic associative search model of recall. The relation between these two independently measured quantities holds even though both of them strongly depend on the presentation rate of the words. We further confirmed the generality of our theory by repeating the experiments with lists of short sentences expressing common knowledge facts. Hence it appears that memory recall is a more universal process than memory acquisition, at least when random material is involved. Since our theory is not specific to the nature of the material being acquired, we conjecture that recall of all other types of information, such as e.g. randomly assembled lists of pictures, should result in similar recall performance.

In the analysis of the model performance, 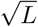 scaling of recall capacity appears when the underlying similarity matrix is assumed to be fully unstructured. This version of the model is equivalent to a random mapping of a finite set to itself which is used to analyze the properties of hashing algorithms in cryptography or the properties of random number generators (Flajolet and Odlyzko [1989]). Here we presented a simple and intuitive analysis of one of the classical statistical properties of random mappings - the average sum of tail and cycle lengths. In contrast, the symmetric similarity matrix model does not have this simple interpretation, since the use of a symmetric matrix imposes non-trivial constraints on mappings, and the rule prohibiting recall of previously recalled words implies a process with memory. Nevertheless, this additional structure still leads to the same *RC* scaling, but with a bigger prefactor. We conjecture that additional specific constraints imposed on SM may lead to increased *RC*, and that human language imposes the set of constraints that lead to dramatic increase of *RC*.

Several influential computational models of recall were developed in cognitive psychology that incorporate interactive probabilistic search processes (see e.g. Raaijmakers and Shiffrin 1980, Gillund and Shiffrin 1984, Howard and Kahana 2002, Laming 2009, Polyn et al. 2009, Lehman and Malmberg 2013). These cognitive models have multiple free parameters that can be tuned to reproduce the experimental results on recall quite precisely, including not only the number of words recalled but also the temporal regularities of recall, such as primacy, recency and temporal contiguity effects (Murdock Jr 1962, Murdock and Okada 1970, Howard and Kahana 1999). However, most of the free parameters lack clear biological meaning and cannot be constrained before the data is collected, hence the models cannot be used to predict the recall performance but only explain it a posteriori. Our recall model can be viewed as a radically simplified version of the classical ‘Search of Associative Memory’ model (SAM), see Raaijmakers and Shiffrin 1980. In both models, recall is triggered by a matrix of associations between the items, which in SAM is built up during presentation according to a rather complex set of processes, while in our model is simply assumed to be a fixed, structure-less symmetric matrix (see Fig. 1). Subsequent recall in SAM proceeds as a series of attempted probabilistic sampling and retrievals of memory items, until a certain limiting number of failed attempts is reached after which recall terminates. In our model, this is replaced by a deterministic transition rule that selects the next item with the strongest association to the currently recalled one. As a result, recall of new items terminates automatically when the algorithm begins to cycle over already recalled items, without a need to any arbitrary stopping rule. Finally, SAM assumes that all the presented words are stored into long-term memory to different degrees, i.e. could in principle be recalled, while in the current study we assume that only a certain fraction of words are effectively acquired to become candidates for recall, the process that we don’t model explicitly but rather access with recognition experiments. We consider it little short of a mystery that with these radical simplifications, the model predicts the recall performance with such a remarkable precision and without the need to tune a single parameter. This suggests that despite all the simplifications, the model faithfully captures a key first-order effect in the data. Future theoretical and experimental studies should be pursued to probe which aspects of the models are valid and which are crucial for the obtained results.

## Acknowledgments

This work is supported by the EU-H2020-FET 1564 and EU-M-GATE 765549 and Foundation Adelis. SR is supported by the Howard Hughes Medical Institute. We thank Drs. Mike Kahana and Eli Nelken for helpful comments.

## Supplemental material

### Methods

#### Recall model

Our recall model is presented in more details in Romani et al. 2013, Katkov et al. 2017. In this contribution we considered a simplified version of the model, where we approximate the matrix of overlaps between random sparse memory representations by a random symmetric *L* by *L* similarity matrix (SM) with otherwise independently distributed elements, where *L* is a number of words in the list. Neglecting the correlations between SM elements is justified in the limit of very sparse encoding of memory items (see Romani et al. 2013). A new matrix is constructed for each recall trial. The sequence {*k*_1_, *k*_2_,…, *k*_*r*_} of recalled items is defined as follows. Item *k*_1_ is chosen randomly among all *L* presented items with equal probability. When *n* items are recalled, the next recalled item *k*_*n*+1_ is the one that has the maximal similarity with the currently recalled item *k*_*n*_, excluding the item that was recalled just before the current one, *k*_*n-*1_. After the same transition between two items is experienced for the second time, the recall is terminated since the model enters into a cycle.

#### Solution of the recall model

The symmetry of SM appears to be a minor difference from the much simpler model of fully random asymmetric SM presented in the main text, but in fact it significantly impacts the statistics of the transitions in the corresponding graphs as we will show below.

If retrieval always proceeds from an item to its most similar, as in the asymmetric case, the dynamics will quickly converge to a two-items loop. The reason is that if item *B* is most similar to item *A*, then item *A* will be most similar to item *B* with a probability of approximately 0.5. We hence let the system choose the second most similar item if the most similar one has just been retrieved, as explained in the main text. When reaching an already visited item, retrieval can either repeat the original trajectory (resulting in a loop) or continue backward along the already visited items and then open a new sub-trajectory (see Fig. 1b). Here we show how to calculate the probability of returning from a new item to any one of already visited items and the probability that the retrieval proceeds along the previous trajectory in the opposite direction upon the return.

In order to return back from item *k* to item *n*, the *n*^*th*^ element of the *k*^*th*^ row of SM, *S*_*kn*_, has to be the largest of the remaining *L* − 2 elements in the *k*^*th*^ row (excluding the diagonal and the element corresponding to the item visited just before the *k*^*th*^ one). The probability for this would be 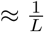 for an asymmetric matrix. For a symmetric matrix (*S*_*nk*_ = *S*_*kn*_), we have an additional constraint that the element *S*_*kn*_ is *not* the largest in the *n*^*th*^ row of *S*, since we require that the *k*^*th*^ item was *not* retrieved after the first retrieval of the *n*^*th*^ one. The probability of return is then equal to

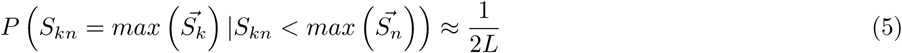

where 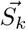 denotes the vector of relevant elements in the *k*^*th*^ row of matrix *S*. The return probability is therefore reduced by a factor of two due to the symmetric nature of SM but retains the same scaling with *L* as in the model with asymmetric SM. After the first return to an item *n* (= 10 in Fig. 1b of the main paper), the trajectory may either begin to cycle, or turn towards previously visited items but in the opposite direction if the original transition from this item (10 → 7 in Fig. 1b) was along the second largest element of 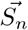. The marginal probability for this is 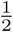, but we must impose the constraint that the *k*^*th*^ item was *not* retrieved after the first retrieval of the *n*^*th*^ one. If the item preceding *n* is *j* (14 in Fig. 1b), the corresponding probability is given by

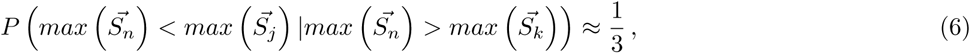

which follows from the observation that any ordering for the maximal elements of three vectors of equal size is equally probable. From this result, we conclude that the average number of sub-trajectories during the retrieval process is 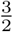. All together the chance for the process to enter a cycle after each new item retrieved is 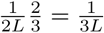 and hence the average number of items recalled is estimated by replacing *L* with 3*L* in the corresponding expression for RC in the model with fully random asymmetric SM, Eq. (2) of the main text:

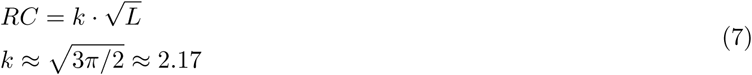

#### Participants, Stimuli and Procedure

In total 1039 participants, were recruited to perform memory experiments on the Amazon Mechanical Turk® (https://www.mturk.com). Ethics approval was obtained by the IRB (Institutional Review Board) of the Weizmann Institute of Science. Each participant accepted an informed consent form before participation and was receiving from 50 to 85 cents for approximately 5 − 25 min, depending on the task. Presented lists were composed of non-repeating words randomly selected from a pool of 751 words produced by selecting English words (Healey et al. 2014) and then maintaining only the words with a frequency per million greater than 10 (Medler and Binder 2005). The stimuli were presented on the standard Amazon Mechanical Turk ® web page for Human Intelligent Task. Each trial was initiated by the participant by pressing “Start Experiment” button on computer screen. List presentation followed 300 ms of white frame. Each word was shown within a frame with black font for 500 or 1000 ms (depending on presentation rate) followed by empty frame for 500 ms. After the last word in the list, there was a 1000 ms delay before participant performed the task. The set of list lengths was: 8, 16, 32, 64, 128, 256 and 512 words. We also performed the same experiments using a set of 325 short sentences expressing well-know facts, such as ‘Earth is round’ or ‘Italians eat pizza’, etc. We repeated our experiments with random lists of 8, 16, 32, 64, and 128 such sentences, each presented for 2500 ms followed by empty frame for 1000 ms. Each participant performed experiment A (free recall) and Experiment B (recognition) with lists of the same length. In more details

- 348 participants performed the two experiments with presentation rate of 1.5 sec/word: 265 participants did both experiments for only one list length, 54 for two list lengths, 18, 9 and 2 for 3, 4 and 5 list lengths respectively.
- 375 participants performed the two experiments with presentation rate of 1 sec/word: 373 participants did both experiments for only one list length, 2 for two list lengths.
- 331 participants performed the two experiments with presentation rate of 3.5 sec/fact: 328 participants did both experiments for only one list length, 3 for two list lengths. 15 participants performed also experiments with the words.

##### Experiment A - Free recall

Participants were instructed to attend closely to the stimuli in preparation for the recalling memory test. After presentation and after clicking a “Start Recall” button, participants were requested to type in as many items (words/sentences) as they could in any order. After the finishing the typing (following non-character input) the information was erased from the screen, such that participants were seeing only the currently typed item. Only one trial was performed by each participant. The time for recalling depended on the length of the learning set, from 1 minute and 30 seconds up to 10 minute and 30 seconds, with a 1 minute and 30 seconds increase for every length doubling. The obvious misspelling errors were corrected. Repetitions and the intrusions (items that were not in the presented list) were ignored during analysis.

##### Experiment B - Recognition task

In recognition trial, after presentation and after clicking a “Start Recognition” button, participants were shown 2 items, one on top of another. One item was randomly selected among just presented in the list (target), and another one was selected from the rest of the pool of words or sentences. The vertical placement of the target was random. Participants were requested to click on the items they think was presented to them during the trial. Each list was followed with 5 recognition trials per participant, but only the first trial was considered in the analysis. Time for all trials was limited to 45 min, but in practice each response usually took less than two seconds.

#### Analysis of the results

The average number of recalled items (*RC*) for each list length and its standard error were obtained from the distribution of the number of recalled items across participants.

In the case of sentences, additional problem that arose concerned the different possible phrasing of the same facts. For example, if a fact presented was ‘Italians like pizza’ and a participant reported ‘Pizza is loved by Italians’, we had to find a way to identify it as correctly recalled. To this end, we used word2vec software developed by Google (Mikolov et al. 2013). Word2vec is a group of related models that are used to produce multidimensional word embeddings. These models are shallow, two-layer neural networks that are trained to reconstruct linguistic contexts of words. Word2vec takes as its input a large corpus of text and produces a vector space, typically of several hundred dimensions, with each unique word in the corpus being assigned a corresponding vector in the space. Word vectors are positioned in the vector space such that words that share common contexts in the corpus are located in close proximity to one another in the space. We used word2vec to compare the sentences reported by participants to the ones presented. A sentence vector was computed as the average of all the word vectors in the sentence, and the similarity between any two sentences with vectors *S*_1_ and *S*_2_ was defined via the cosine of the angle between them, as 1 − cos (*S*_1_, *S*_2_). If the similarity was greater than 0.9 the recall was considered to be correct (this threshold was confirmed by manual inspections of multiple cases). The cases in which the similarity was between 0.8 and 0.9 were checked manually.

The average number of items acquired for each list of length *L* was computed from the results of recognition experiments as in (Standing 1973). Suppose that *M* out of *L* items are remembered on average after an exposure to the list, the rest are missed. The chance that one of the acquired items is presented during a recognition trial is then *M/L*, while the chance that a missed word is presented is 1 − *M/L*. We assume that in the first case, a participant correctly points to a target item, while in the second case, she/he is guessing. The fraction of correct responses *c* can then be computed as

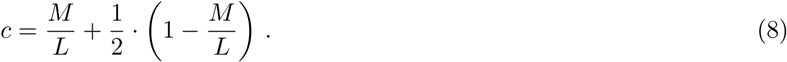

Hence the average number of remembered items can be computed as

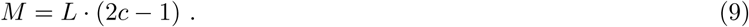

In order to estimate a standard error of the mean for the number of acquired items across participants, for each list length, we performed a bootstrap procedure (Efron and Tibshirani 1994). We generated multiple bootstrap samples by randomly sampling a list of N participants with replacement N times. Each bootstrap sample differs from the original list in that some participants are included several times while others are missing. For each bootstrap sample *b* out of total number *B*, with *B* = 500, we compute the estimate for the average number of acquired items, *M* (*b*), according to Eq. (9). The standard error of *M* is then calculated as a sample standard deviation of *B* values of *M* (*b*):

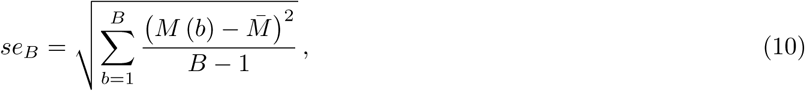

where 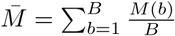

## Additional figures

**Figure S1.**
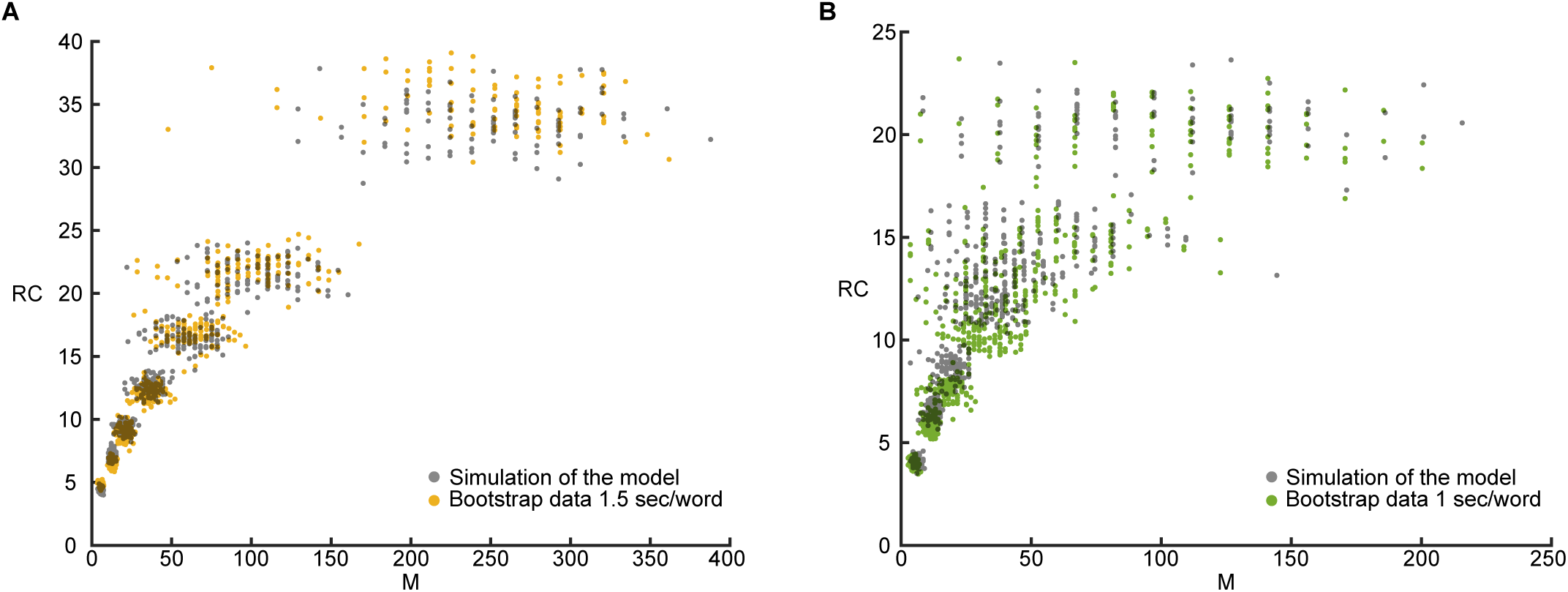
Bootstrap analysis and comparison to model simulations. 1.5 seconds per word presentation rate; (**B**) 1 second per word presentation rate. 100 bootstrap samples for each list length are shown with colored dots with coordinates *M* (*b*) and *RC*(*b*), where *RC*(*b*) is an average number of recalled words computed for each bootstrap sample *b*. Black dots show corresponding simulation results, obtained as follows. From the results of recognition experiment, we calculate, for each list length *L*, the fraction of correct recognitions across the participants, *c*, and therefore the probability *p* = (2*c* − 1) that a presented word is acquired. With these two numbers, we simulate multiple recognition and recall experiments. For recognition experiment, we draw a binomial random variable with probability *c* for each participant independently, simulating their recognition answers, from which we compute the number of acquired words averaged for all participants as explained in the Methods. We then drew *L* binomial variables with probability *p* for each participant, simulating the acquisition of words by this participant during the recall experiment. With the number of acquired words known for each participant, we run the recall model (see Methods) to obtain the average recall performance over participants. Every simulation described above produced 7 pairs of results (*M, RC*), one per list length. We repeated the whole procedure 100 times, same as the number of bootstrap samples.

